# AUG-3387, a Human-Derived Monoclonal Antibody Neutralizes SARS-CoV-2 Variants and Reduces Viral Load from Therapeutic Treatment of Hamsters In Vivo

**DOI:** 10.1101/2021.10.12.464150

**Authors:** Christopher J. Emig, Marco A. Mena, Steven J. Henry, Adela Vitug, Christian John Ventura, Douglas Fox, Xammy Huu Nguyenla, Haiyue Xu, Chaeho Moon, Sawittree Sahakijjpijarn, Philip J. Kuehl, David Revelli, Zengrong Cui, Robert O. Williams, Dale J. Christensen

**Affiliations:** Augmenta Bioworks, 3475 Edison Way, Suite K, Menlo Park, CA 94025 USA; School of Public Health, Division of Infectious Diseases and Vaccinology, University of California, Berkeley, 1951 Oxford Street, Berkeley, CA 94720; Department of Molecular and Cellular Biology, University of California, Berkeley, 1951 Oxford Street, Berkeley, CA 94720; Molecular Pharmaceutics and Drug Delivery Division, College of Pharmacy, The University of Texas at Austin, 2409 University Avenue, Austin, TX, 78712, USA; TFF Pharmaceuticals, Inc., Austin, TX, 78746 USA; Lovelace Biomedical Research Institute, Albuquerque, NM 87108, USA

**Keywords:** Pulmonary administration, Thin-Film Freezing, monoclonal antibody, Dry powder inhaler, SARS-CoV-2 therapy, hamster model

## Abstract

Infections from the SARS-CoV-2 virus have killed over 4.6 million people since it began spreading through human populations in late 2019. In order to develop a therapeutic or prophylactic antibody to help mitigate the effects of the pandemic, a human monoclonal antibody (mAb) that binds to the SARS-CoV-2 spike protein was isolated from a convalescent patient following recovery from COVID-19 disease. This mAb, designated AUG-3387, demonstrates a high affinity for the spike protein of the original viral strains and all variants tested to date. *In vitro* pseudovirus neutralization and SARS-CoV-2 neutralization activity has been demonstrated in vitro. In addition, a dry powder formulation has been prepared using a Thin-Film Freezing (TFF) process that exhibited a fine particle fraction (FPF) of 50.95 ± 7.69% and a mass median aerodynamic diameter (MMAD) and geometric standard deviation (GSD) of 3.74 ± 0.73 µm and 2.73 ± 0.20, respectively. The dry powder is suitable for delivery directly to the lungs of infected patients using a dry powder inhaler device. Importantly, AUG-3387, administered as a liquid by intraperitoneal injection or the dry powder formulation delivered intratracheally into Syrian hamsters 24 hours after intranasal SARS-CoV-2 infection, demonstrated a dose-dependent reduction in the lung viral load of the virus. These data suggest that AUG-3387 formulated as a dry powder demonstrates potential to treat COVID-19.

## 1. Introduction

Infections caused by a novel coronavirus began emerging in late 2019 that became known as the Severe Acute Respiratory Syndrome Coronavirus 2 (SARS-CoV-2) virus. This Coronavirus Disease 2019 (COVID-19) pandemic has resulted in the loss of more than 4.6 million lives globally to date. The SARS-CoV-2 virus is related to the etiologial agents causing the SARS epidemic in 2002-2003 and the Middle East Respiratory Syndrome (MERS) epidemic of 2012 (*1, 2*). The COVID-19 pandemic has resulted in more deaths than any other recent epidemics (*1, 3, 4*) and has had profound effects on the lives of people around the world. One key mitigation strategy employs neutralizing monoclonal antibodies (mAbs) to treat or protect against SARS-CoV-2 infection. Since the outbreak, multiple mAb products have been granted Emergency Use Authorization (EUA) but all are administered by infusion or injection. These include Casirivimab and Imdevimab produced by Regeneron, Bamlanivimab and Etesevimab marketed by Lilly, and Sotrovimab from Glaxo Smith Klein in partnership with Vir Biologics. Although generally limited to infusion centers and out-patient settings, mAb therapies have proven to reduce hospitalization in mild to moderate COVID-19 patients.

Routes for discovery of new SARS-CoV-2 neutralizing antibodies include isolation of antibody sequences from patients who have recovered from SARS-CoV-2 or SARS-CoV-1, inoculation and isolation of humanized mice, or the use of phage or other library display technology. Over the past two decades, several groups have published methods for isolating, sequencing and cloning antibody genes from single B cells from primary patient samples, then expressing antibody protein for characterization (*5-7*). We have developed workflows for discovery of antigen specific membrane bound antibodies via cytometry, and secreted antigen-specific antibodies via our proprietary SingleCyte^®^ instrument (Augmenta Bioworks). SingleCyte^®^ is a motorized microscope with an intelligent image processing engine driving a single cell retrieval device. A typical SingleCyte^®^ assay measures antigen binding of secreted antibodies against 4 antigen targets for up to 240,000 single cells in a standard SBS format plate and retrieves approximately 200 antigen specific single B cells (Figure 1). Single cells isolated using SingleCyte or FACS are deposited into 96 well plates and heavy and light chain mRNAs are reverse transcribed, amplified and sequenced. The sequences are then PCR amplified or synthesized and cloned into expression vectors. We have developed a fully integrated and highly automated platform for performing these workflows in under ten days and have had previous success in identifying antibodies targeting other infectious disease antigens, including herpes simplex virus and influenza virus.

**Figure 1.**
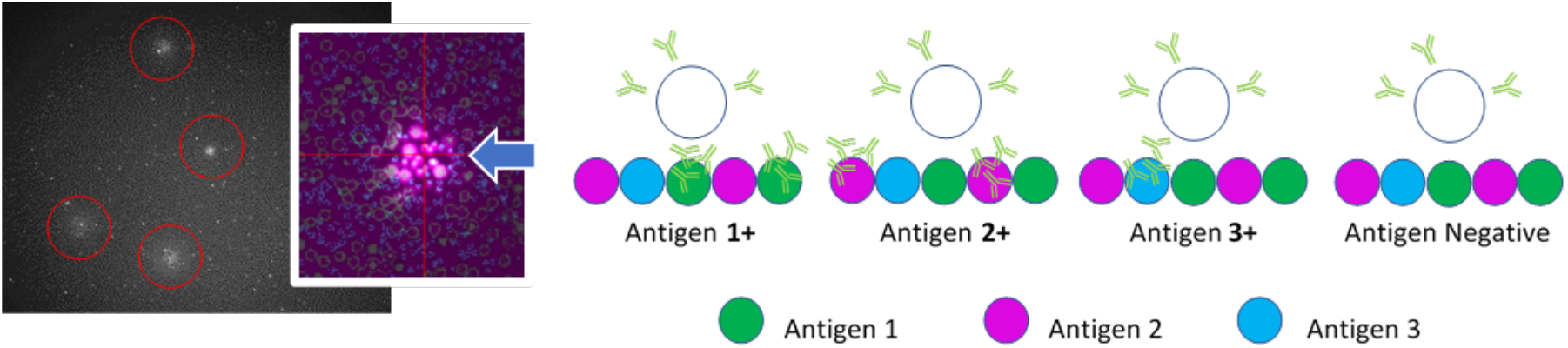
Illustration of the SingleCyte^®^ cell isolation process. Cells are assayed for their ability to secrete proteins that bind various optically encoded antigens in their proximity and target cells are selected and isolated using an automated system.

Since SARS-CoV-2 is primarily a pulmonary disease, early treatment or prophylaxis with neutralizing mAb therapy targeted to the airways could improve disease outcomes. Initiating treatment early in the disease cycle may alter the course of the disease, prevent the development of chronic complications, reduce the hospitalization rate and decrease mortality. Only a fraction of a systemically administered mAb is transported into the pulmonary compartment where viral particles are released early in the disease; therefore, delivery of neutralizing mAbs directly to the lung holds the potential to reduce the dose needed to achieve the same efficacy as systemically delivered mAbs. Aerosolized delivery would have additional benefits including the potential for patients to self-administer antibody therapy at home rather than in infusion centers, and the ability to expand the supply of antibody to treat a larger population through dose-reduction. Furthermore, a dry antibody formulation stable at ambient temperatures could be made available to patients in geographic regions that lack suitable infrastructure for cold chain distribution of injectible antibody formulations that require cold chain distribution.

Thin-film freezing (TFF) is a particle engineering technology that has been used to prepare dry powder formulations of drugs that are administered to patients using a dry powder inhaler (DPI) device. Powders produced by this process are currently in human clinical testing for the treatment of pulmonary indications (NCT04872231 and NCT04576325) (*8, 9*) and have characteristic highly porous brittle matrices with low bulk densities that can be delivered with good aerosol performance (*10*). The TFF technology generates these powders by fast supercooling of drug-carrier solutions (*11*) followed by lyophilization to remove water or other solvents and has been used to generate powders of small molecule (*12, 13*) and biologic drugs (*14*).

We now report the isolation of a fully human mAb, AUG-3387, using the SingleCyte^®^ system, which binds to the SARS-CoV-2 spike protein with high affinity and binds to all SARS-CoV-2 variant spike proteins tested to date including Delta, Lamda and Mu. In vitro neutralization was confirmed using pseudovirus neutralization with the wild-type SARS-CoV-2 and the Delta variant (B.1.617.2) spike protein pseudotyped lentivirus. Direct neutralization of wild-type SARS-CoV-2 was also demonstrated in vitro where AUG-3387 demonstrated a 99.99% reduction of infection of VERO-E6 cells at therapeutically relevant doses, which demonstrated that AUG-3387 has potential to treat or prevent COVID-19 disease. In order to translate these findings to prepare for clinical trials, AUG-3387 was formulated into a dry powder formulation that is stable at ambient temperatures using the Thin Film Freezing process as a powder that contains 15% weight to weight of the antibody. The dry powder was delivered to Syrian golden hamsters in an in vivo therapeutic model of COVID-19 disease where treatment with AUG-3387 was initiated 24 hr after inoculation with SARS-CoV-2. In this model, three daily intratracheal insufflation administrations of the dry powder or a single intraperitoneal injection of AUG-3387 in liquid formulation demonstrated a dose-responsive reduction in viral titer in lung tissue at day 5. Taken together, these data demonstrate the potential for AUG-3387 to treat or prevent COVID-19.

## 2. Materials and Methods

### 2.1. Materials

The reagents and suppliers listed in Table 1 were used to complete the studies.

**Table 1:** Reagents and Supplies.

| Reagent | Supplier | Catalog Number |
| --- | --- | --- |
| SARS-CoV-2 Wuhan-1 RBD | Sino Biological | 40592-V08B |
| B.1.1.17 (Alpha) S1 | Sino Biological | 40591-V08H12 |
| 20H/501Y.V2 (Beta) S1 | Sino Biological | 40591-V08H10 |
| B.1.617.2 (Delta) RBD | Sino Biological | 40592-V08H86 |
| B.1.1.28 (Gamma) S1+S2 | Sino Biological | 40589-V08B10 |
| B.1.617 (Kappa) RBD | Sino Biological | 40592-V08H88 |
| SARS-CoV-2 S494P RBD | Sino Biological | 40592-v08h18 |
| SARS-CoV-2 V483A RBD | Sino Biological | 40592-v08h5 |
| SARS-CoV-2 R683A, R685A, F817P, A892P, A899P, A942P, K986P, V987P S1+S2 | Sino Biological | 40589-v08h4 |
| SARS-CoV-2 G485S RBD | Sino Biological | 40592-v08h52 |
| SARS-CoV-2 D614G,E484K S1 | The Native Antigen Company | REC31902 |
| SARS-CoV-2 D614G, V445I, H655Y, E583D S1 | The Native Antigen Company | REC31903 |
| SARS-CoV-2 Spike-E-M Mosaic Protein | The Native Antigen Company | REC31829-100 |
| SARS-CoV-2 (2019-nCoV) Spike Protein (S2 ECD, His tag) | Sino Biological | 40590-V08B |
| SARS-CoV-2 Spike Glycoprotein (S1) "Supplier 2" | The Native Antigen Company | REC31828-100 |
| SARS-CoV-2 (2019-nCoV) Spike Protein (S1+S2 ECD, His tag) | Sino Biological | 40589-V08B1 |
| SARS-CoV-2 (2019-nCoV) Spike Protein (S1 Subunit, His Tag) "Supplier 1" | Sino Biological | 40591-V08H |
| SARS-CoV-2 L452R,T478K RBD | Sino Biological | 40592-V08H90 |
| SARS-CoV-2 Lenti Pseudovirus | GenScript | SC1993-2 |
| SARS-CoV-2 Delta Lenti Pseudovirus | e-Enzyme | SCV2-PsV-Delta |
| BSA | ROCKLAND IMMUNOCHEMICALS | BSA-50 |
| HEK293T | ATCC | CRL-3216 |
| pCMV-AC-GFP Ace2 Expression Vector | Origene | pCMV-AC-GFP |
| ACE2 Polyclonal antibody | ProteinTech | 21115-1-AP |
| PE Donkey anti-rabbit IgG | BioLegend | 406421 |
| MagPlex -C Microspheres | Luminex | MC10012-01, etc (various) |
| Antibody Conjugation Kit | Luminex | 40-50016 |
| Thermo Scientific Zeba Spin Desalting Columns, 7K MWCO, 0.5 mL | Thermo Scientific | PI89883 |
| 1 xMAP Sheath Fluid, 20L, RUO | Luminex | 40-50015 |
| PE anti-human IgG/IgA/IgM | Sigma | AQ503H |
| Thermo Scientific Zeba Spin Desalting Columns, 7K MWCO, 0.5 mL | Thermo Scientific | PI89883 |
| Anti-V5 antibody clone SV5-Pk1 | Abcam | ab27671 |
| BrightGlo Luciferase Reagent | Promega | E2620 |

### 2.2. Antibody Isolation

For this study, we profiled patients with low disease burden, rather than severely affected individuals. We hypothesized that asymptomatic or weakly symptomatic patients might have had previous exposure to a related antigen (e.g., other coronaviruses), providing a breadth of antigenic coverage and driving their resistance to COVID-19. Informed consent was obtained from all patients, and all patient samples were collected after a full recovery from illness and under IRB approval. Patient samples were profiled to identify samples with binding to multiple SARS-CoV-2 proteins including the S1-receptor binding domain (RBD), full length S1, S2, E and M protein. From each sample, plasma was separated from PBMC’s and red blood cells by Ficoll density gradient centrifugation (Cytiva). Additionally, approximately 100k B cells were extracted from whole blood (RosetteSep, StemCell) and seeded directly, or terminally differentiated into plasma cells and then seeded into SingleCyte^®^ assay plates with up to 4 different optically encoded SARS-CoV-2 antigens or controls. A fluorescently labelled anti-human IgG/IgA/IgM secondary antibody was also added to each well. After a 24 hour incubation, antigen specific cells were identified by their signature reaction-diffusion pattern. For identification of surface bound antibodies binding to SARS-CoV-2, SARS-CoV-2 antigens conjugated to Alexa Fluor 488 stained SARS-CoV-2 antigen was used as a staining agent for single cell sorting of antigen reactive memory B cells into 96 well plates on a Sony SH-800 Cell Sorter. Cells specific to any SARS-CoV or SARS-CoV-2 antigen were retrieved and sequenced.

The sequences were analyzed for unique clonotypes based on CDR3 sequences of heavy and light chains, then unique clonotypes were designed into DNA fragments for cloning through Augmenta’s AbWorks™ automated clone design software. The fragments were cloned as ScFv’s for expression in E. coli, or as full heavy and light chains in mammalian expression vectors for tandem transfection into Expi293T.

SingleCyte^®^ is Augmenta’s programmable single cell imaging cytometer and sorter that selects cells based on temporal microscopy. For screening of secreted antibody proteins, assay plates contain a multiplexed panel of antigens in the form of conjugated beads or antigen presenting cells and a secondary antibody in solution. Antibodies from secreted cells bind proximal antigens and become physically constrained near the secreting cell. Fluorescent secondary antibody enables visualization of secreted antibody concentration gradients based on fluorescence over time, and optically encoded antigen beads enables deconvolution of target antigens. Assays are performed in standard open well SBS footprint microplates and are user programmable. The instrument works by first raster imaging each well. Cells with any antigen reactivity are identified by processing images with a convolutional neural network trained on a set of manually curated images. The microscope performs multispectral high resolution imaging of each positive cell. Images are masked into regions by the optical characteristics of proximal beads and a confidence score is ascribed to each cell-antigen interaction. Single cells are then picked and placed into receiver plates and post-isolation images are taken to ensure proper aspiration of target cells. A robotic arm carries receiver plates for high throughput single cell retrieval. The information for each run and output metrics for each cell (including both source and destination locations) are saved to a database for recall in a user interface and for downstream processing steps.

### 2.3. Antibody Chracterization

#### 2.3.1 Target Binding Characterization

SARS-CoV-2 S1 and RBD proteins, and various other antigens and controls, were covalently coupled to Luminex MagPlex magnetic microspheres for assay binding with a Luminex 200 instrument. The following antigens were conjugated: B.1.1.17 (Alpha) S1, B.1.1.28 (Gamma) S1+S2, 20H/501Y.V2 (Beta) S1, B.1.617 (Kappa) RBD, B.1.617.2 (Delta) RBD, S1+S2 S494P, S1+S2 V483A, S1+S2 R683A+R685A+F817P+A892P+A899P+A942P+ K986P+V987P, S1+S2 G485S, S1+S2 D614G, S1+S2 E484K, S1+S2 D614G+V445I+H655Y+E583D, S1+S2 L452R+T478K. Each antigen was conjugated with the xMAP conjugation kit at ratio of 5µg protein to 1 million beads. Assays were performed in multiplex, with each spectrally encoded bead having a separate antigen and run together in a single well. Antibody was titrated over therapeutically relevant concentrations, mixed with the beads, washed twice, labelled with a secondary antibody, washed twice and run on the instrument. Dry powder versions of antibodies were resuspended in water before dilution for assay.

The ScFv version of AUG-3387, designated AUG-3705, was run on a Carterra LSA instrument at multiple concentrations for determination of the single domain affinity against Wuhan SARS-CoV-2 S1 and RBD. AUG-3705 was attached to the LSA flow cell via interaction with its V5 tag and a surface bound anti-V5 antibody. Wuhan-1 RBD was delivered to the flow cell at concentrations of 2.06nM, 6.17nM, 18.5nM and 56nM for calculation of Kd.

### 2.4. Pseudoneutralization Assay

#### 2.4.1 ACE-2 Expressing HEK293T Cell Line Construction

An ACE2 expressing HEK293T cell line (“LentiX ACE2.S4”) was constructed by packaging pCMV-AC-GFP (Origene) into lentivirus and transducing HEK293T’s (ATCC). The cells were enriched 4 times until 97% of the cells showed signal above the negative control as read out by staining with anti-ACE-2 and secondary antibodies. On average enriched ACE2-HEK293T’s had 50-fold higher signal compared to the signal of non-transduced cells.

#### 2.4.2 SARS-CoV-2 Pseudovirus with TFF powder formulation and soluble AUG-3387

Two days prior to infection, LentiX ACE2.S4 cells were grown to 85% confluency, then seeded in a 96-well plate at 15k cells/well in 50µL media per well and held at 37 °C in 5% CO_2_ until infection. Antibody mixes were created prior to infection by performing a 128-fold serial dilution starting at 40µg/µL. Dry powders prepared by the TFF process of AUG-3387 and soluble negative control V5 Tag monoclonal antibody were seeded in triplicate, and soluble AUG-3387 was seeded in duplicate. SARS-CoV-2 pseudovirus (Genscript) was diluted in DMEM complete media to an IFU of 3.2e7/mL, and 100µL of virus solution was mixed with 100 µL of diluted antibody. The virus/antibody mix was incubated for 60 minutes at 37 °C in 5% CO_2_. Following incubation, 50 µL of each pseudovirus/antibody condition mix was added to each well of seeded cells. Additional controls included cells only, and cells with virus only. After 48 hours, the plate was removed and equilibrated at room temperature for 10 minutes, and 60 µL of the supernatant was removed. 50 µL of Promega’s Bright-Glo Luciferase assay reagent was added to each well of the infected cells. The cells then were incubated at room temperature for 3 minutes, and luminescence was measured with a Tecan Spark microplate reader with a 1 second integration time.

#### 2.4.3 SARS-CoV-2 Pseudovirus, Delta Variant (B.1.617.2) with soluble AUG-3387

Two days prior to infection, LentiX ACE2.S4 cells were grown to 85% confluency, then seeded in a 96-well plate at 15k cells/well in 50µL media per well and held at 37°C in 5% CO_2_ until infection. Antibody mixes were created prior to infection by performing a 128-fold serial dilution starting at 160µg/µL. AUG-3387 and negative control V5 Tag monoclonal antibody were seeded in triplicate. SARS-CoV-2 Delta Variant pseudovirus (eEnzyme) was diluted 1:2 in DMEM complete media to a pseudoviral particle concentration of 5e7/ml, and 200µL of virus solution was mixed with 200µL of diluted antibody. The virus/antibody mix was incubated for 60 minutes at 37 °C in 5% CO_2_. Following incubation, 100µL of each pseudovirus/antibody condition mix was added to each well of seeded cells. Additional controls included cells only, and cells with virus only. After 48 hours, the plate was removed and equilibrated at room temperature for 10 minutes, and 100µL of the supernatant was removed. 50µL of Promega’s Bright-Glo Luciferase assay reagent was added to each well of the infected cells. The cells then were incubated at room temperature for 3 minutes, and luminescence was measured with a Tecan Spark microplate reader with a 1 second integration time.

### 2.5. SARS-CoV-2 Neutralization Assay

Two days prior to infection, Calu-3 cells were grown to confluency, then seeded at 40k cells in 100µL media per well. Antibody was titrated in D10 media. The reagents were then transferred into a Bio Safety Level 3 facility for further processing. For each antibody condition, SARS-CoV-2 virus at a target MOI of 0.05 was mixed with antibody at the desired concentration and incubated at 37 °C for 60 minutes. Media was removed from the seeded cells and replaced with a final volume of 50 µL of antibody/virus mix. Cells with antibody and virus (MOI 0.05) were incubated at 37 °C in 5 % CO2 for 30 minutes. The virus/antibody mix was removed and the cells were washed with 75 µL PBS. Cells were replenished with titrated antibody in a final volume of 150 µL media.

After 24 hours, 50 µL of the supernatant was removed for TCID50 assays. Vero E6 cells were seeded at 10,000 cells in 100 µL per well. Infected cell culture supernatant was diluted with 950 µL D10 media, and then serial diluted before 50 µL of each dilution was added to 8 wells of Vero E6 cells. After 72 hours, wells with complete cytopathic effect were counted. After 96 hours, 100 µL CellTiterGlo reagent was added to each well of the infected Calu-3 cells to assay for live cells. Following incubation of CTG reagent for 20 minutes, luminescence was measured with a Spectramax 1L with 1s integration time.

### 2.6. Preparation of Thin Film Freezing (TFF) composition

In the preparation of the solutions for TFF manufacturing, AUG-3387 was combined with a mannitol/leucine or trehalose/leucine. The solution was applied as drops onto a rotating cryogenically cooled drum cooled to -70 °C. The frozen solids were collected and stored in a -80 °C freezer before lyophilization. The lyophilization was performed in an SP VirTis Advantage Pro shelf lyophilizer (SP Industries, Inc., Warminster, PA, USA). The primary drying process was at -40 °C for 20 h, and then, the temperature was linearly increased to 25 °C over 20 h, followed by secondary drying at 25 °C for 20 h. The pressure was maintained at less than 100 mTorr during the lyophilization process.

### 2.7. Aerodynamic particle size distribution analysis

About three milligrams of AUG-3387 mAb dry powder was loaded into size #3 hydroxypropyl methylcellulose (HPMC) capsules (Vcaps^®^ plus, Capsugel^®^, Lonza, Morristown, NJ, USA). The aerodynamic properties of the powder were evaluated using a Next Generation Impactor (NGI) (MSP Corporation, Shoreview, MN, USA) connected to a High-Capacity Pump (model HCP5, Copley Scientific, Nottingham, UK) and a Critical Flow Controller (model TPK 2000, Copley Scientific, Nottingham, UK). A high-resistance Plastiape^®^ RS00 inhaler (Plastiape S.p.A, Osnago, Italy) was used for dispersing the powder through the USP induction port with a total flow rate of 60 L/min for 4 s per each actuation corresponding to a 4 kPa pressure drop across the device and a total flow volume of 4 L. To avoid particle bounce, a solution of polysorbate 20 in methanol at 1.5% (w/v) was applied and dried onto the NGI collection plates to coat their surface. The pre-separator was not used in this analysis. After dispersal, the powder was extracted from the stages using water.

Quantitation of mannitol or trehalose recovered from NGIstages was performed on an Agilent 1220 Infinity II HPLC system (Santa Clara, CA) with a Waters XBridge Amide column (4.6 × 150 mm, 3.5 µm) (Milford, MA) connected to Agilent 1290 Infinity II ELSD (Santa Clara, CA). A gradient method with mobile phase B, from 80% to 40% acetonitrile with 0.1 % (v/v) trifluoroacetic acid, and mobile phase A, water, was used at the mobile phase flow rate of 1.0 mL/min and the column temperature of 30°C. 15 µL of each sample was injected and ran for 6 minutes. Evaporative and nebulizer temperatures of ELSD were set at 60°C, and the gas (dry nitrogen) flow rate was 1.6 L/min.

The analysis was conducted three times (n=3). The NGI results were analyzed using the Copley Inhaler Testing Data Analysis Software 3.10 (CITDAS) (Copley Scientific, Nottingham, UK). CITDAS provided the calculation for mass median aerodynamic diameter (MMAD), geometric standard deviation (GSD), fine particle fraction (FPF) of delivered dose (FPF%, delivered) and recovered dose (FPF%, recovered). The FPF of delivered dose was calculated as the total amount of sugar or sugar alcohol (e.g., trehaose, mannitol) collected with an aerodynamic diameter below 5 µm as a percentage of the total amount of sugar or sugar alcohol deposited on the adapter, the induction port, stages 1–7 and Micro-Orifice Collector.

### 2.8. Efficacy of AUG-3387 in and in vivo model of SARS-CoV-2 infected Syrian Hamsters

An *in vivo* efficacy study was performed with male Syrian Hamsters (*Mesocricetus auratus*) approximately 9 weeks of age with a weight range of 110-134 g, at time of randomization, were sourced from Charles River Laboratory. Animal work was performed at Lovelace Biomedical Research Institute (LBRI), with approval from the Institutional Animal Care and Use Committee (IACUC) and within Animal Biosafety Level 3 (ABSL3) containment. Hamsters were singly housed in filter-topped cage systems and were supplied with a certified diet, filtered municipal water, and dietary and environmental enrichment. The challenge study design is detailed in Table 2. Animals were assigned to groups using a stratified (body weight) randomization procedure. Animals were anesthetized and swabs of nasal passages were collected by placing the nasal swab (0.5 mm diameter Ultrafine Micro Plasdent swabs) 1-3 mm into the nare and swabbing. Lung and nasal swab (in Trizol) samples were stored at -80°C prior to analysis. All animals were euthanized with an euthanasia solution consisting of 390 mg of sodium pentobarbital and 50 mg of phenytoin per mL.

**Table 2:** Group Designations of animals in the efficacy study.

| Group | Group Description | Challenge Day | Dose | Route/Frequency | Number of animals | Study Endpoints |
| --- | --- | --- | --- | --- | --- | --- |
| 1 | Vehicle Control | Day 0 | 0.0 | IP; once on Day 1 | 6 | Daily Clinical Observations<br>Twice Daily<br>Body Weights Daily<br>Nasal swabs on Days 1 and 5<br>for viral load<br>Necropsy on day 5 for viral<br>load of lung tissue and<br>Histopathology of lungs |
| 2 | mAb-1 High dose | Day 0 | 1.0 mg/kg | IT; once daily on Days 1,2,3 | 6 |  |
| 3 | mAb-1 Low dose | Day 0 | 0.33 mg/kg | IT; once daily on Days 1,2,3 | 6 |  |
| 4 | mAb-solution High dose | Day 0 | 10.0 mg/kg | IP; once on Day 1 | 6 |  |
| 5 | mAb-solution Low dose | Day 0 | 3.3 mg/kg | IP; once on Day 1 | 6 |  |

#### 2.8.1 Viral Challenge

SARS-CoV-2, isolate USA-WA1/2020, was sourced from WRCEVA and propagated in Vero E6 African Green Monkey kidney cells (BEI, catalog #N596) in Dulbecco’s Modified Eagle Medium supplemented with 1% HEPES, 10% FBS, 100 IU/mL Penicillin G and 100 μg/mL Streptomycin. Stocks were stored in a BSL-3 compliant facility at -80°C prior to challenge. Stock vials of virus were thawed the day of challenge, diluted as necessary, and stored on wet ice until use. Viral challenge dose was quantitated using a Tissue Culture Infectious Dose 50% (TCID_50_) assay using the Reed and Muench method (*15*) on Vero E6 cells in DMEM supplemented with 2% FBS and 100 IU/mL Penicillin G and 100 μg/mL Streptomycin. A challenge dose of 1.0 × 10^5^ TCID_50_ per animal was targeted. Actual challenge dose averaged 5.8 × 10^5^ TCID_50_ per animal. The viral challenge dose was delivered via intranasal installation under anaesthesia (ketamine 80 mg per kg and xylazine 5 mg per kg) with a volume of 100 µL per nare (200 µL total per animal).

#### 2.8.2 AUG-3387 Treatment

The AUG-3387 mAb was devliered by one of two routes for each animal, IT and IP injection. The IP injection was performed with a 16 mg/mL formulation in saline. Intratracheal insufflation was performed with animals under anesthesia (4-5% isoflurane with oxygen) until a deep plane of anesthesia was reached. Dry powder for inhalation delivery was transferred to the ABSL-3 facility and each individual device quantitiatively loaded for delivery. Doses were based on method development to quantify the amount of material that exited the devices assuming 100% presentation at the terminus of the canuale and the animals average body weight during dosing.

#### 2.8.3 Quantitative Assessment of Viral Burden

Quantitation of genomic viral RNA and subgenomic viral RNA, by RT-qPCR was used as markers for viral burden. Nasal swab and lung samples were assayed via RT-qPCR for both the N-gene (genomic) and the E-gene (subgenomic). For both methods, lung samples were weighed and homogenized using a Tissue Lyser (Qiagen) in 1 ml of TRI reagent. RNA was extracted using the Direct-Zol 96-RNA kit (Zymo Research) according to manufacturer’s instructions. RNA was quantified using qRT-PCR TaqMan Fast Virus 1-step assay (Applied Biosystems). SARS-CoV-2 specific primers and probes from the 2019-nCoV RUO Assay kit (Integrated DNA Technologies) were used: (L Primer:TTACAAACATTGGCCGCAAA; R primer: GCGCGACATTCCGAAGAA; probe: 6FAM-ACAATTTGCCCCCAGCGCTTCAG-BHQ-1). Reactions were carried out on a BioRad CFX384 Touch instrument according to the manufacturer’s specifications. A semi-logarithmic standard curve of synthesized SARS-CoV-2 N gene RNA (LBRI) was obtained by plotting the Ct values against the logarithm of cDNA concentration and used to calculate SARS-CoV-2 N gene in copies per gram of tissue or per nasal swab.

Copies of SARS-CoV-2 E gene were measured by qRT-PCR TaqMan Fast Virus 1-step assay (Thermo Fisher). SARS-CoV-2 specific primers and probes from the 2019-nCoV RUO Assay kit (Integrated DNA Technologies) were used: (L Primer: CGATCTCTTGTAGATCTGTTCTC; R primer: ATATTGCAGCAGTACGCACACA; probe: 6FAM-ACACTAGCCATCCTTACTGCGCTTCG-BHQ-1). Reactions were carried out on a BioRad CFX384 Touch instrument according to the manufacturer’s specifications. A semi-logarithmic standard curve of synthesized SARS-CoV-2 E gene RNA (LBRI) was obtained by plotting the Ct values against the logarithm of cDNA concentration and used to calculate SARS-CoV-2 E gene in copies per gram of tissue or per nasal swab. Thermal cycling conditions involved 5 minutes at 50°C for reverse transcription, followed by an initial denaturation step for 20 seconds at 95°C and 40 cycles of 95°C for 3 seconds and 60°C for 30 seconds.

## 3. Results and Discussion

### 3.1. Antibody Isolation

In order to isolate antibodies that bind and neutralize SARS-CoV-2, samples from seven study subjects were profiled for the presence of SARS-CoV-2 antibodies using a Luminex-based profiling 6-fold multiplex assay (Supplemental Figure 1). A proprietary instrumentation platform known as SingleCyte^®^ was used to select SARS-CoV-2-reactive antibody-producing B-cells. SingleCyte^®^ is a programmable single cell imaging cytometer and sorter that selects cells based on temporal microscopy. For screening of secreted antibody proteins, assay plates contain a multiplexed panel of antigens in the form of conjugated beads or antigen presenting cells and a secondary antibody in solution. Antibodies from secreted cells bind proximal antigens and become physically constrained near the secreting cell. Fluorescent secondary antibody enables visualization of secreted antibody concentration gradients based on fluorescence over time, and optically encoded antigen beads enables deconvolution of target antigens (Figure 2a). Assays are performed in standard open well SBS footprint microplates and are user programmable. A custom nanoliter volume micropipette enables isolation of single cells, and a robotic arm carries receiver plates for high throughput single cell retrieval.

**Figure 2:**
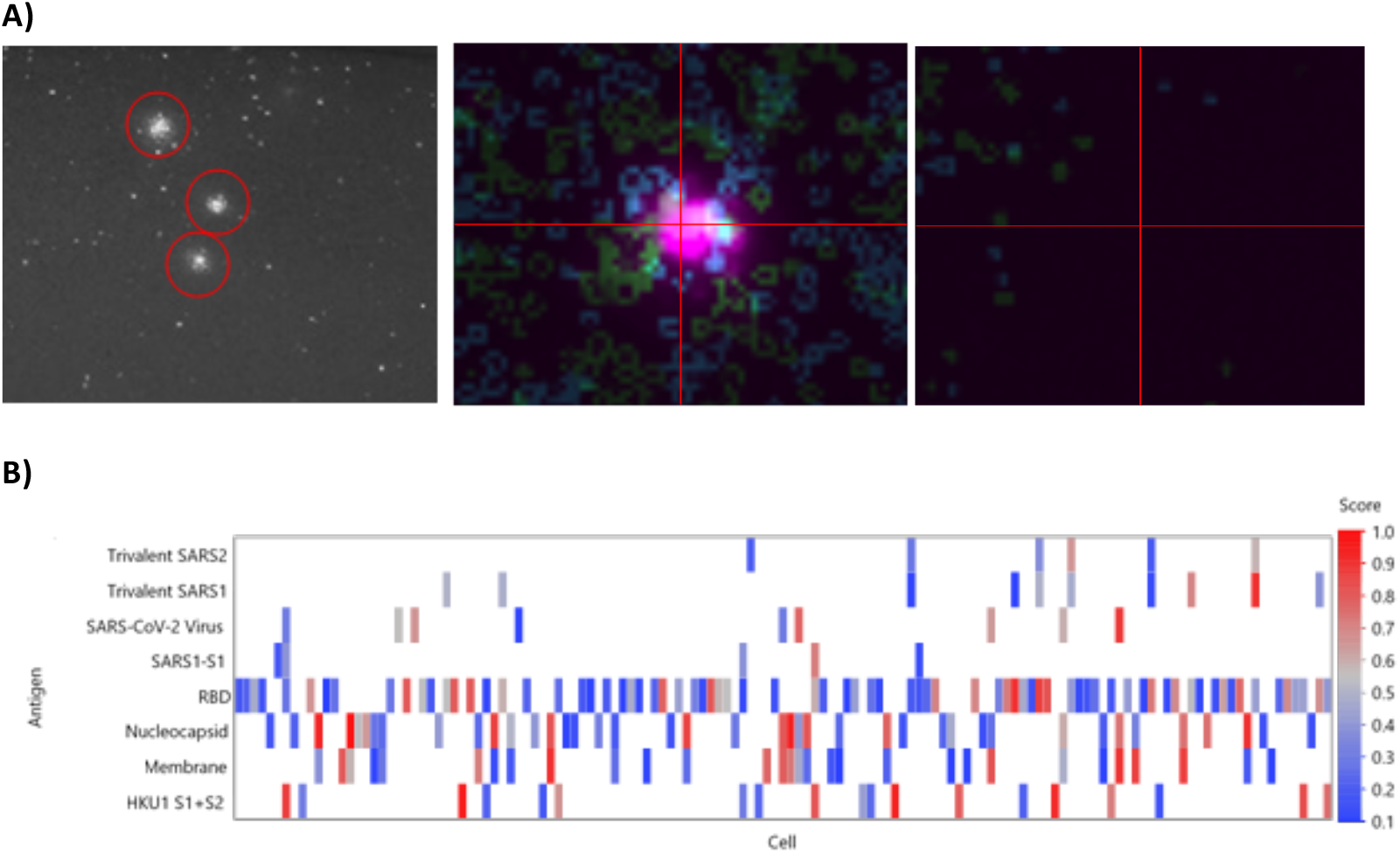
Isolation and Characterization of SARS-CoV-2 binidng Cells: A) Raster of plate at 5X showing antigen specific B cells with signature reaction-diffusion pattern. Before and after 20X false color images of automated capture of an S1-RBD specific plasma cell. Blue indicates antigen beads displaying S1 RBD protein, while green indicates beads displaying S2 protein. Magenta is a cell specific stain. B) Example output from a SingleCyte Screen. For each cell, the confidence of an antigen specific interaction is determined by the amount of secondary antibody signal that overlaps an antigen-specific bead image in the proximity of each cell. A score of 0 represents a 50% chance of antigen specificity, with 1.0 representing a 100% certainty.

Approximately 800 single cells were isolated with SingleCyte and 200 with single cell flow sorting. From these cells, nearly 500 paired chain antibody constructs were designed and approximately 200 were expressed and assayed (Figure 2b). We recovered many S1, S2, and RBD binders and ultimately chose AUG-3387 as our lead compound due to its breadth of binding activity, affinity to the Wuhan-1 strain, and strength in viral neutralization. The timeline for isolation process is shown in Supplemental Figure 2.

### 3.2. Antibody Characterization

#### 3.2.1 AUG-3387 Single Domain Affinity

We utilized the Carterra LSA platform to determine the single domain affinity of AUG-3387 expressed as an ScFv, designated AUG-3705. Auto-fitting of curves was performed in Carterra Kinetics software, which returned a calculated affinity of 1.2 nM (Fig 3).

**Figure 3.**
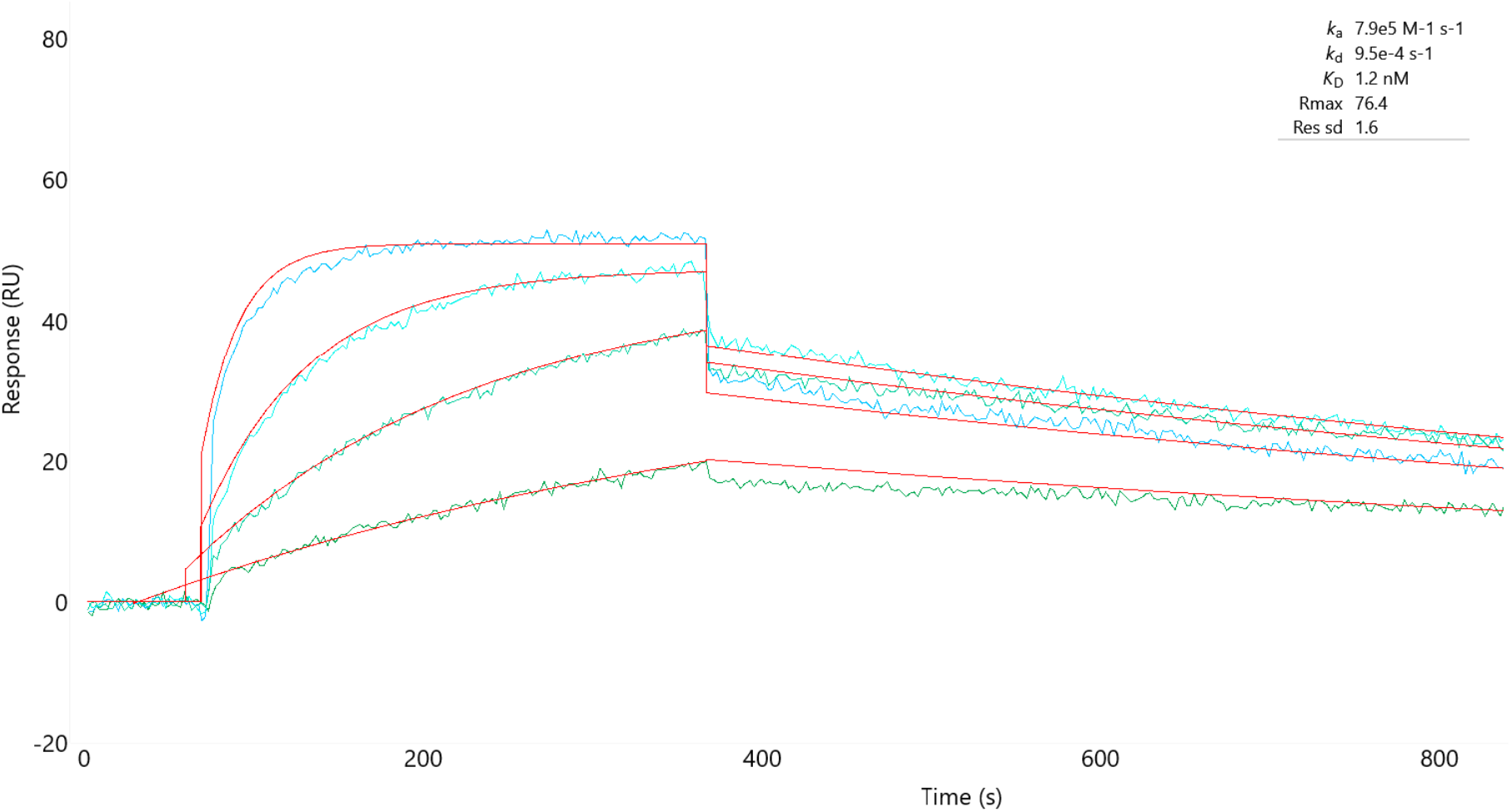
Single domain affinity of AUG-3387 binding domain expressed as an ScFv. The Carterra LSA platform was used to measure affinity of the single chain version of AUG-3378 by flowing increasing concentrations of SARS-CoV-2 spike protein and measuring binding by surface plamon resonance.

#### 3.2.2 AUG-3387 Variant Binding Assays

To assess the susceptibility of AUG-3387 to mutational escape, we profiled AUG-3387 against the S1 and RBD portions of the original Wuhan-1 strain of SARS-CoV-2, RBD’s corresponding to WHO designated dominant strains of concern, and S1 mutants known to affect the potency of currently approved therapeutic antibodies. AUG-3387 binds every S1 and RBD of SARS-CoV-2 tested with a binding EC50<200ng/ml (Figure 4), but only very weakly to SARS-CoV-1 (binding EC50 > 100ug/ml, not shown). These data demonstrate the high potential for resistance to mutational escape of AUG-3387.

**Figure 4:**
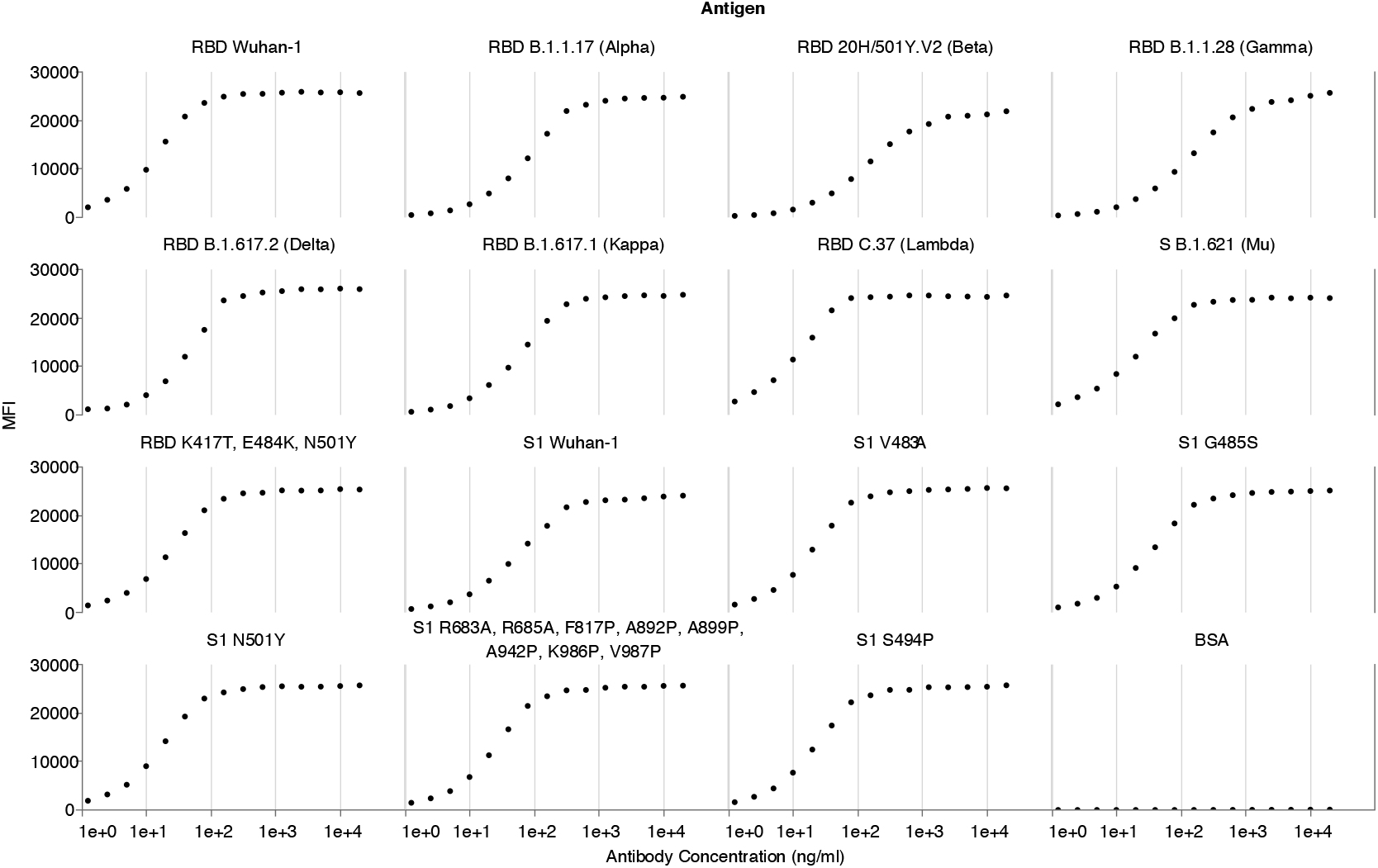
illustrates the Resistance of AUG-3387 to Mutational Escape. AUG-3387 was profiled against the S1 and RBD portions of the original Wuhan-1 strain of SARS-CoV-2, RBD’s corresponding to WHO designated dominant strains of concern, and S1 mutants known to affect the potency of currently approved therapeutic antibodies. AUG-3387 binds every S1 and RBD of SARS-CoV-2 tested with a binding EC_50_ <200ng/ml, but only very weakly to SARS-CoV-1 (binding EC_50_ > 100ug/ml, not shown).

#### 3.2.3 AUG-3387 Neutralization of SARS-CoV-2 Wuhan-1

We compared the ability of full length IgG1 formatted AUG-3387 and its ScFv formatted version, AUG-3705, to neutralize live SARS-CoV-2 in a 24 hour TCID50 assay and a 96 hour infected cell viability assay (Figure 5). AUG-3705 demonstrated somewhat higher efficacy in these assays over AUG-3387, indicating the improved avidity of the dimeric IgG1 did not improve neutralization enough to compensate for the higher molarity of AUG-3705 at the same concentration.

**Figure 5.**
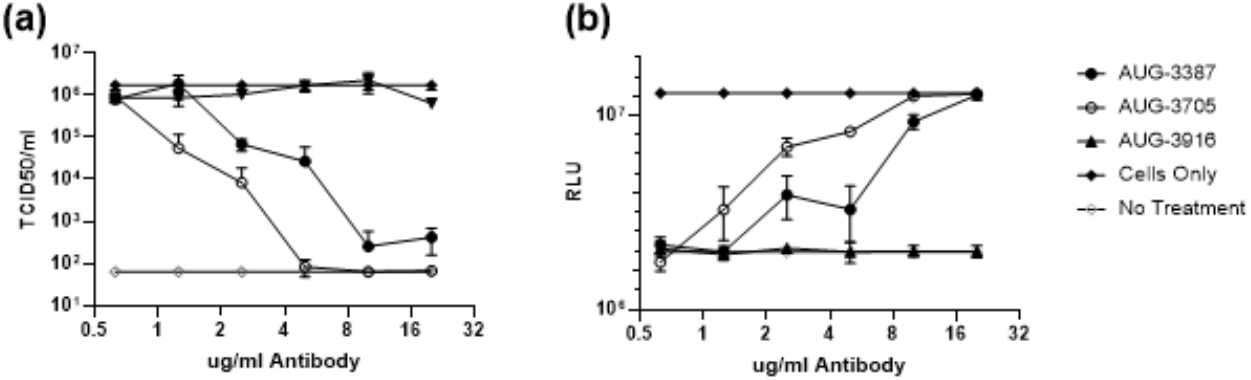
Neutralization of SARS-CoV-2 by AUG-3387 and AUG-3705. Neutralization was determined by measuring TCID50 24 hours post infection (a) and measuring for viable cells (CellTiterGlo luminescence) 96 hours post infection (b). In both assays, the dose-response curves indicate an IC50 of ∼2ug/ml or less.

#### 3.2.3 AUG-3387 Neutralization of Delta Pseudovirus

We assessed the ability of AUG-3387 to neutralize the SARS-0CoV-2 Delta variant pseudotyped virus (Figure 6). AUG-3387 demonstrated the ability to neutralize Delta pseudovirus, although at a higher IC50 (30-40 mg/mL) than for the Wuhan-1 strain (approximately 2 mg/mL in TCID50 assay). However, the 30-40 mg/mL is still a clinically relevant dose that can be achieved by delivery to the lung.

**Figure 6.**
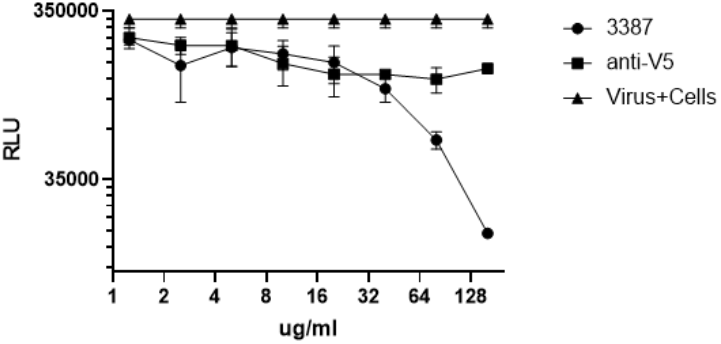
Neutralization of SARS-CoV-2 B.1.617.2 (Delta) pseudovirus by AUG-3387. Neutralization was determined by measuring luciferase activity of cells infected with Delta pseudovirus after treatment with AUG-3387.

### 3.3. TFF Powder Optimization and Characterization

A series of dry powder formulations of AUG-3387 were prepared by Thin-Film Freezing and evaluated for retention of biological activity and optimal aerosol properties for delivery to the lung. The powders contained AUG-3387 at a range of mAb concentrations from 5-20% (w/w) and various excipients. The powders were tested for the presences or absence of subvisible aggregates under a light microscope, for mAb aggregation or fragmentation using SDS-PAGE, and for their aerosol properties using the NGI. We assessed two formulations of AUG-3387 in TFF powders, AUG-3387.11 prepared with mannitol/leucine (95%/5%) and AUG-3387.13 prepared with trehalose/leucine (95%/5%), and compared them to the original AUG-3387 formulation in PBS. AUG-3387.11 and AUG-3387.13 performed virtually identically to their soluble counterpart in gel electrophoresis (Figure 7A), multiplexed bead assays (Figure 7B).

**Figure 7.**
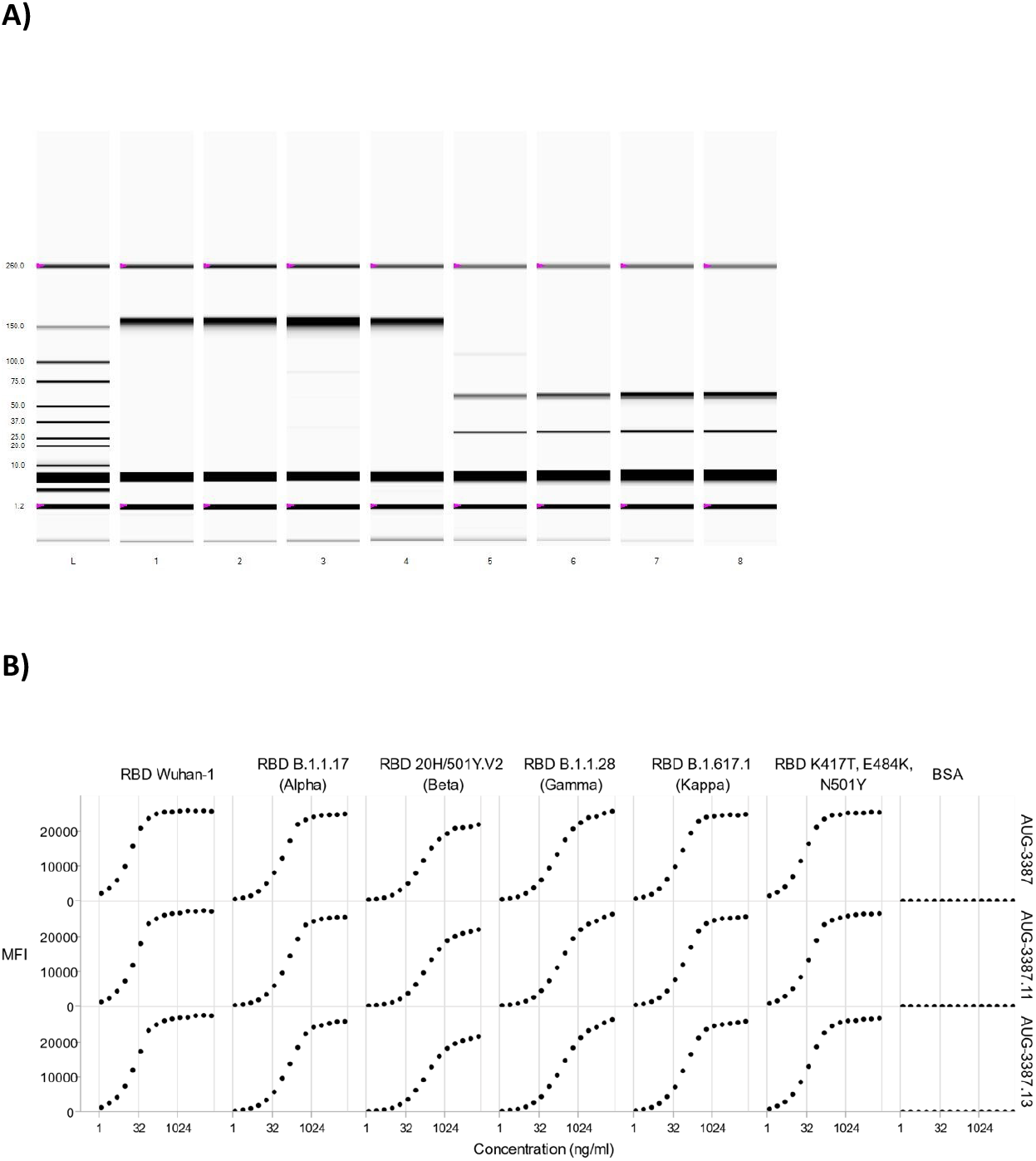
Characterization of Dry Powder Formulations of AUG-3387 Prepared by Thin Film Freezing. A) Samples were run on a Bio-Rad Experion with AUG-3387.11 and AUG-3387.13 native in lanes 1 and 2, reducing in lanes 5 and 6, respectively. AUG-3387 input mAb as expressed in Expi-293T in lane 3 and 7 and CHO in lane 4 and 8. B) AUG-3387.11 and AUG-3387.13 bind to SARS-CoV-2 variants at the same concentrations as the PBS formulation indicating that no loss of binding occurred after the TFF processing to create dry powder formulations.

After verification that the dry powder formulations demonstrated no loss of binding activity, a pseudoneutralization assay was used to confirm that the biological activity was retained. The dry powder formulations were dissolved in media and added to the pseudoneutralization assay along with the input antibody. Both dry powder formulations, AUG-3387.11 and AUG-3387.13, retained full neutralization activity of the parental mAb (Figure 8).

**Figure 8.**
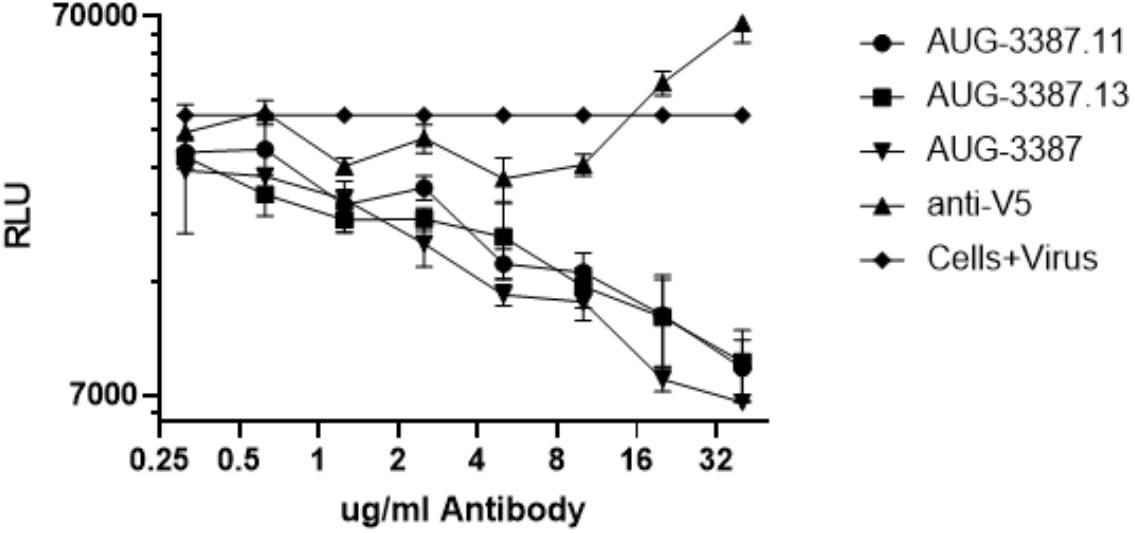
Pseudoneutralization activity of Dry Powder Formulated AUG-3387. and dry powder formulations AUG-3387.11 and AUG-3387.13 demonstrate neutralization of SARS-CoV-2 Wuhan-1 pseudovirus at the same concentration as the PBS formulation

Final selection of the AUG-3387 dry powder formulation for *in vivo* characterization was complete by evaluating the aerosol properties of the powders. The dry powder designated AUG-3387.11 showed excellent aerosol properties (Figure 9). This powder, delivered through a Plastiape RS00 Dry powder inhaler, gave an MMAD value of 3.74 ± 0.73 µm, a GSD of 2.73 ± 0.20, and an FPF (delivered) of 50.95 ± 7.69%. Upon reconstitution of the powder, no significant subvisible aggregated particles in the solution were observed under a microscope.

**Figure 9.**
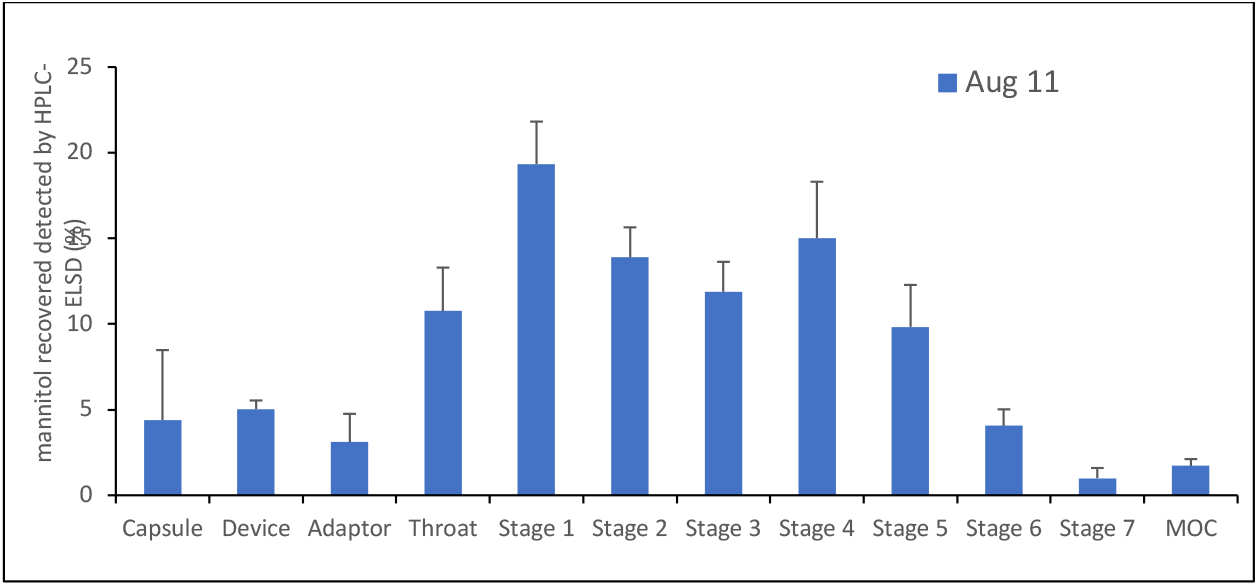
NGI Characterization of AUG-3387.11. Aerodynamic particle size distribution of AUG-3387.11 was determined by actuating a capsule for delivery using RS00 high-resistence DPI at a flow rate of 60 L/min (n =3) and quantitation of the powder on each stage of the impactor using an HPLC-ELSD method.

### 3.4. AUG-3387 Reduces SARS-CoV-2 Viral Load in Syrian Golden Hamsters

The efficacy of AUG-3387 for therapeutic reduction of viral load was assessed *in vivo* using the established hamster model with the mAb formulations being delivered starting 24 hours after intranasal SARS-CoV-2 inoculation. Hamsters were administered AUG-3387 at doses of 3 and 10 mg/kg or a vehicle control by intraperitoneal (IP) injection. Additional groups received three doses of the TFF dry powder formulation of AUG-3387 by intratracheal (IT) instillation of at doses of 0.3 and 1 mg/kg at 24, 48, and 72 hours after SARS-CoV-2 inoculation. All animals showed body weight loss. On Day 5, animals were harvested and lung tissues were assessed for viral replication by rt-qPCR for subgenomic (active) viral replication. Dose dependent viral load reductions were observed with both the IP and IT treated animals showing reduced viral load in the lung tissue, despite treatment not being initiated until 24 hours after intranasal SARS-CoV-2 inoculation.

**Figure 10.**
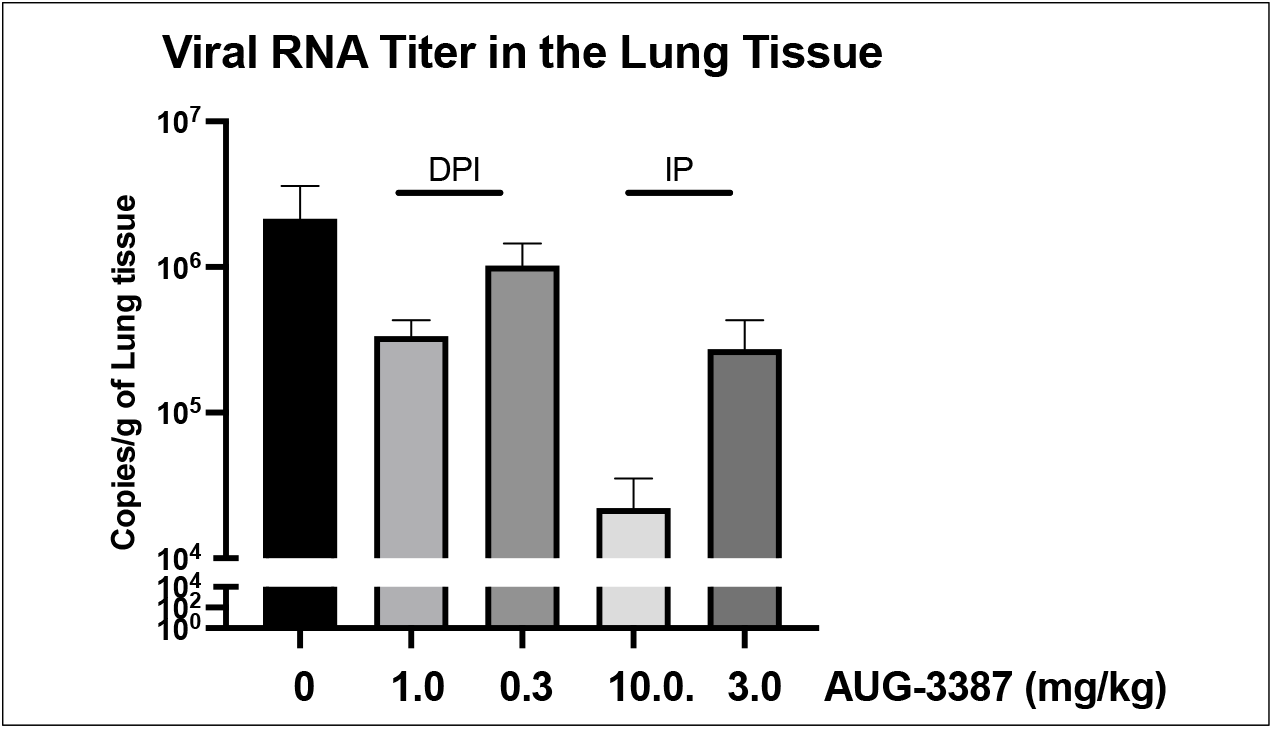
Viral Load of SARS-CoV-2 Virus in Hamster Lung Tissue. Following inoculation with SARS-CoV-2 at time 0, treatement was initated 24 hr later and aninmals were euthanized on Day 5. Lung tissue was homogenized and active viral load was characterized by RT-qPCR for the E-gene.

## 4. Discussion

Since the start of the COVID-19 disease pandemic caused by the SARS-CoV-2, a high burden has been placed on the healthcare system to provide adequate care and treatment of the high number of patients infected by the virus that require hospitalization. Patient care has improved as the pandemic has progressed and the mortality rate has dropped as the care has improved. One key area of improvement in patient outcomes occurred when anti-SARS-CoV-2 monoclonal antibody therapies became available. While not highly efficacious for the treatment of severe COVID-19 disease, the current mAb therapies demonstrate up to a 79% reduction in hospitalization in patients that are symptomatic and at high risk to develop severe disease that will require hospitalization. These risk factors include age greater than 65 years, obesity or being overweight, pregnancy, diabetes, chronic kidney disease, immunocompromised patients due to disease or immunosuppressive treatment, cardiovascular disease, chronic lung diseases or sickle cell disease.

In these patients, mAb therapy has been demonstrated to reduce hospitalization by up to nearly 80% when administered early in the course of disease when the patients present with mild to moderate disease.

However, each of the mAb products currently authorized under EUA by the FDA are administered by intravenous infusion, which requires the patient of have the infusion performed in a healthcare setting where infusion personnel are adequately trained to perfom intravenous catheter placement and administration of the IV solutions. While these mAb therapeutics provide a clear therapeutic benefit for the reduction of hospitalization, the route of administration continues to place a burden on the healthcare system to deliver these therapies to the patients and their families. Thus, delivery of mAb therapy to patients in an outpatient setting without the need for specialized infusion centers would provide a clear advancement. Furthermore, since only a fraction of systemically administered antibodies reaches the lung from systemic administration, localized delivery to the lung would represent an advantage over intravenous delivery because the virus replicates in the pulmonary epithelial cells early in the infection and can be administered at lower total doses per patient.

The SingleCyte^®^ system was used to isolate a new mAb, designated AUG-3387, that displays potent binding to the SARS-CoV-2 S-protein. Binding to and neutralization of both pseudovirus and Wuhan-1 Coronavirus demonstrated the potential utility of AUG-3387 for treatment of COVID-19 disease. AUG-3387 demonstrated potent binding to the Alpha, Beta, Gamma, Delta, Lamda and Mu variants suggesting that AUG-3387 binds a conserved epitope that has notmutated in the variants of concern or newly emerging Lamda and Mu variants. Furthermore, neutralization and pseudoneutralization data demonstrate that AUG-3387 prevents the virus from infecting cells by blocking interaction of this conserved region of the RBD with the hACE2 receptor of target cells. The retained activity against all tested variants is in contrast to the reduced susceptibility of Bamlanivimab and Etesevimab, which show greater than 250-fold reduced binding and neutralization activity against the Beta and Gamma variants (*16*).

In order to differentiate AUG-3387 from the current mAb therapeutics that are currently being used under EUA, the mAb has been formulated as a room temperature stable dry powder utilizing the thin-film freezing process. The room temperature stability may allow for distribution to geographic locations where SARS-CoV-2 continues to spread but that do not have the capability of distributing injectible formulations that require cold chain distribution and storage. We demonstrated that the formulations prepared using the TFF process retain full binding activity of the input mAb solutions and have no evidence of protein aggregation or instability following reconstitution in water.

In addition to the dry powder storage at room temperature of the TFF formulated dry powder mAb, the dry powders can be encapsulated and delivered to the lung using a standard dry powder inhaler device. When tested with the Plastiape RS00 high resistance device, which is designed to provide maximum shear and aerosolization of powders at lower airflow rates, the AUG-3387 powder formulations had a fine particle fraction with greater than 50% of the powder in the 1-5 μm range, which is ideal for delivery to the deep lung of humans using a device matched to the potential for reduced lung function for mild to moderate COVID-19 patients.

Finally, we demonstrated that administration of AUG-3387 by either intraperitoneal injection or by intratracheal insufflation of the dry powder into SARS-CoV-2 infected Syrian hamsters resulted in a dose dependent reduction of the viral load in the lung tissue of the infected hamsters. The result of this *in vivo* study is notable because we utilized a treatment paradigm that creates a high burden for efficacy to be demonstrated. In our study, mAb treatment of the hamsters was not initiated until 24 hours after the hamsters were infected with SARS-CoV-2 by intranasal inoculation. By contrast, sotrovimab administered by IP injection prophylactically at doses of 5 mg/kg or more when given 24- or 48-hours prior to viral infection resulted in improvement in body weight loss and decreased viral load in the lung tissue compared to control animals. Likewise, the casirivimab and imdevimab combination of mAbs administered to hamsters by IP injection 24 hours before viral inoculation (*17*) resulted in a dose dependent viral load reduction in lung tissue. However, no change in viral load in the lung tissue was reported when casirivimab and imdevimab were administered 24 hours after viral inoculation in a manner similar to our study. For sotrovimab there was no report of therapeutic treatment resulting in reduced viral load. Thus, to date, the demonstration that AUG-3387 administered by either IP or IT routes in a therapeutic mode resulted in a dose dependent viral load reduction in the lung tissue represents the first report of a mAb therapy that works in the hamster model in a therapeutic mode. Furthermore, the viral load reduction of the dry powder when delivered by IT insufflation represents the first report of successful reduction of viral load using inhaled delivery of a mAb therapeutic for COVID-19 disease.

Taken together, these data suggest that AUG-3387 is a potent mAb that has the potential to treat all known variants of SARS-CoV-2 and that the powders produced by the TFF formulation process to make room temperature stable powders has the potential to reduce the amount of mAb needed for efficacy because of the local delivery to the lung. Furthermore, since the TFF AUG-3387 powder does not require cold chain storage, it represents an opportunity to distribute the powder formulation globally to reduce the human cost of the COVID-19 pandemic by facilitating delivery of this therapy to locations lacking cold chain distribution capabilities.

## Acknowledgements

The authors would like to thank the laboratory of Peter S. Kim for sharing trimeric SARS-CoV1/2 spike proteins and performing preliminary pseudoneutralization assays, Sarah A. Stanley for access to BSL3 facilities at UC Berkeley, and Daniel Bedinger of Carterra LSA for determining affinity of AUG-3705 and other Augmenta antibodies. We also greatly appreciate the contributions to Augmenta’s platform and discovery efforts by Payam Shahi, John Beaber, Rosanna Chau, Cindy Y. Lai, Kim X. Nguyen, Robin Emig, John R. Haliburton, Max Von Franque, Niklas Mannhardt, Isaac Perper, Micah Kelly, Sarah Galloway, Rachel Brewer, Paige Hansen, Jeremiah Budiman, Helen Chang, and William Horvat.

---

**Supplemental Figure 1.**
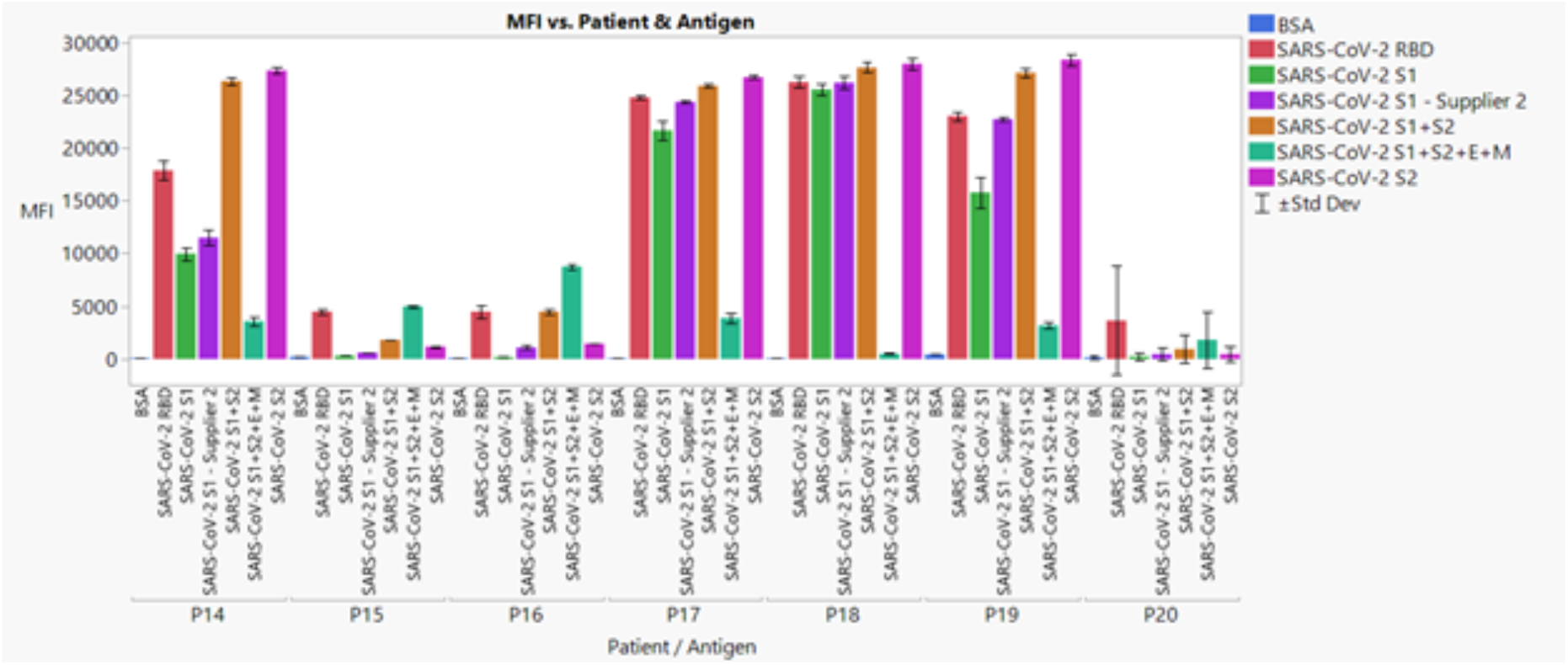
Serological profile of 7 patients against 6 SARS-CoV-2 antigens generated by diluting plasma 1:1000 in a 6 plex Luminex assay. Samples from each patient were evaluated for binding to antigen proteins including the Receptor binding domain (RBD), Spike Protein 1 (S1), Spike Protein 2 (S2), the E-protein (E), and M protein (M) coated on Luminex beads.

**Supplemental Figure 2.**
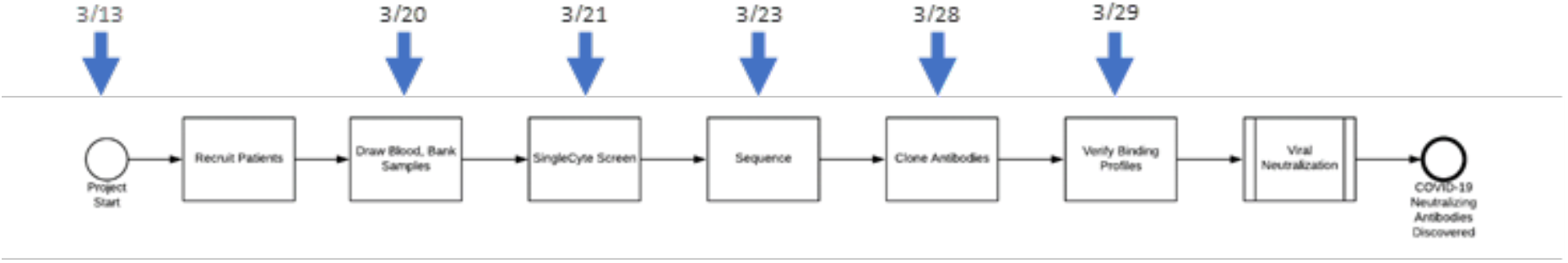
Timeline for discovery of Augmenta’s first SARS-CoV-2 antibodies.

## References

1. K. Miller, M. E. McGrath, Z. Hu, S. Ariannejad, S. Weston, M. Frieman, W. T. Jackson, Coronavirus interactions with the cellular autophagy machinery. Autophagy 16, 2131–2139 (2020).

2. F. Wu, S. Zhao, B. Yu, Y.-M. Chen, W. Wang, Z.-G. Song, Y. Hu, Z.-W. Tao, J.-H. Tian, Y.-Y. Pei, A new coronavirus associated with human respiratory disease in China. Nature 579, 265–269 (2020).

3. D. Gurwitz, Angiotensin receptor blockers as tentative SARS-CoV-2 therapeutics. Drug development research 81, 537–540 (2020).

4. G. Xue, X. Gan, Z. Wu, D. Xie, Y. Xiong, L. Hua, B. Zhou, N. Zhou, J. Xiang, J. Li, Novel serological biomarkers for inflammation in predicting disease severity in patients with COVID-19. International immunopharmacology 89, 107065 (2020).

5. H.-X. Liao, M. C. Levesque, A. Nagel, A. Dixon, R. Zhang, E. Walter, R. Parks, J. Whitesides, D. J. Marshall, K.-K. Hwang, High-throughput isolation of immunoglobulin genes from single human B cells and expression as monoclonal antibodies. Journal of virological methods 158, 171–179 (2009).

6. T. Tiller, E. Meffre, S. Yurasov, M. Tsuiji, M. C. Nussenzweig, H. Wardemann, Efficient generation of monoclonal antibodies from single human B cells by single cell RT-PCR and expression vector cloning. Journal of immunological methods 329, 112–124 (2008).

7. J. Wrammert, K. Smith, J. Miller, W. A. Langley, K. Kokko, C. Larsen, N.-Y. Zheng, I. Mays, L. Garman, C. Helms, Rapid cloning of high-affinity human monoclonal antibodies against influenza virus. Nature 453, 667–671 (2008).

8. U.S. National Library of Medicine. (2021, May 6). Identifier NCT04872231, Single Ascending Dose and Multiple Ascending Dose Study of Voriconazole Inhalation Powder in Healthy Adult Subjects. Clinicaltrials.Gov. https://clinicaltrials.gov/ct2/show/NCT04872231

9. U.S. National Library of Medicine. (2021, October 6). Identifier NCT04576325, Pharmacokinetic Profile of Voriconazole Inhalation Powder in Adult Subjects With Asthma. Clinicaltrials.Gov. https://clinicaltrials.gov/ct2/show/NCT04576325

10. N. A. Beinborn, J. Du, N. P. Wiederhold, H. D. Smyth, R. O. Williams, 3rd, Dry powder insufflation of crystalline and amorphous voriconazole formulations produced by thin film freezing to mice. Eur J Pharm Biopharm 81, 600–608 (2012).

11. C. Moon, S. Sahakijpijarn, J. J. Koleng, R. O. Williams, Processing design space is critical for voriconazole nanoaggregates for dry powder inhalation produced by thin film freezing. Journal of Drug Delivery Science and Technology 54, (2019).

12. C. Moon, A. B. Watts, X. Lu, Y. Su, R. O. Williams, 3rd, Enhanced Aerosolization of High Potency Nanoaggregates of Voriconazole by Dry Powder Inhalation. Mol Pharm 16, 1799–1812 (2019).

13. S. Sahakijpijarn, C. Moon, X. Ma, Y. Su, J. J. Koleng, A. Dolocan, R. O. Williams, 3rd, Using thin film freezing to minimize excipients in inhalable tacrolimus dry powder formulations. Int J Pharm 586, 119490 (2020).

14. S. Hufnagel, S. Sahakijpijarn, C. Moon, Z. Cui, R. O. Williams III, The Development of Thin-film Freezing and Its Application to Improve Delivery of Biologics as Dry Powder Aerosols. KONA Powder and Particle Journal, 2022010 (2022).

15. L. J. Reed, H. Muench, A simple method of estimating fifty per cent endpoints. American journal of epidemiology 27, 493–497 (1938).

16. Fact sheet for health care providers emergency use authorization (EUA) of bamlanivimab and etesevimab. (2021, September 16). U.S. FDA. https://www.fda.gov/media/145802/download

17. A. Baum, D. Ajithdoss, R. Copin, A. Zhou, K. Lanza, N. Negron, M. Ni, Y. Wei, K. Mohammadi, B. Musser, REGN-COV2 antibodies prevent and treat SARS-CoV-2 infection in rhesus macaques and hamsters. Science 370, 1110–1115 (2020).

